# Revealing Age-Related Changes of Adult Hippocampal Neurogenesis

**DOI:** 10.1101/112128

**Authors:** Frederik Ziebell, Sascha Dehler, Ana Martin-Villalba, Marciniak-Czochra Anna

## Abstract

In the adult hippocampus, neural stem cells (NSCs) continuously produce new neurons that integrate into the neuronal network to modulate learning and memory. The amount and quality of newly generated neurons decline with age, which can be counteracted by increasing intrinsic Wnt activity in NSCs. However, the precise cellular changes underlying this age-related decline or its rescue through Wnt remain unclear. The present study combines development of a mathematical model and experimental data to address features controlling stem cell dynamics. We show that available experimental data fit a model in which quiescent NSCs can either become activated to divide or undergo depletion events, consisting of astrocytic transformation and apoptosis. Additionally, we demonstrate that aged NSCs remain longer in quiescence and have a higher probability to become re-activated versus being depleted. Finally, our model explains that high NSC-Wnt activity leads to longer time in quiescence while augmenting the probability of activation.

## 1 Introduction

The subgranular zone of the hippocampal dentate gyrus (DG) is one of the two major regions in the adult brain where neural stem cells (NSCs) continuously produce new neurons involved in the process of learning and memory [1]. The age-associated impairment of learning and memory has awaken much interest in understanding the cellular and molecular mechanisms underlying the accompanying decline of NSC numbers in the hippocampus [2, 3, 4, 5, 6, 7]. Hence, interpretations of the dynamics of NSCs in homeostasis have led to many recent studies [8, 9, 10, 11, 12, 13].

To decipher the cellular dynamics behind hippocampal neurogenesis, we apply an interdisciplinary approach combining mathematical modeling with experimental data. These data consist of already published studies [11, 8, 9, 12, 6] as well as novel data specifically designed to reveal age-related changes during hippocampal neurogenesis. Our approach of combining modeling with data was successfully applied for other stem cell-based systems [14, 15, 16, 17, 18], in which mathematical models were used to identify and quantify stem cell-related processes such as self-renewal and differentiation.

The developed mathematical model describes the evolution of the hierarchical cell production system of adult hippocampal neurogenesis and incorporates basic cell properties such as quiescence, proliferation, self-renewal and apoptosis. Since the exact regulatory feedbacks in neurogenesis are not well understood, we follow a top-down approach in which we start with a basic model with constant rates and then apply it to experimental data in order to identify which parameters are changing during aging.

Our study shows that the data are consistent with a model in which NSCs reside in a quiescent phase from which they can either become activated to proliferate or undergo depletion events. By applying our model to novel data displaying an age-related accumulation of astrocyte numbers, we demonstrate that about 40% of NSC depletion can be attributed to direct transformation into astrocytes with the remaining part being likely a result of apoptosis.

Finally, we use our model to uncover the response of the neurogenesis system upon Dickkopf-1 (Dkk1) deletion [6]. We show that the best explanation for the effects upon Dkk1 knockout is that stem cells spend longer time in quiescence but are also more likely to become re-activated versus being depleted from the quiescent phase.

## 2 Results

### 2.1. Stem Cell Model Fits Population-level and Clonal Data

Our model is based on a compartment of quiescent NSCs, corresponding to the *G*_0_ phase of the cell cycle (Figure 1). Those quiescent NSCs have the ability to enter the cell cycle and perform a symmetric self-renewing or asymmetric division, followed by NSCs returning to quiescence after division [8]. Although Bonaguidi et al. [8] do not address a mechanism leading to NSC disappearance, they find an increased number of NSC-depleted stem cell clones in their data and conclude that there are NSC depletion events. Accordingly, we assume an outflow from the quiescent NSCs compartment not only due to cell cycle entry but also due to depletion. As shown later, our results suggest that NSC depletion consists of apoptosis as well as astrocytic transformation.

**Figure 1:**
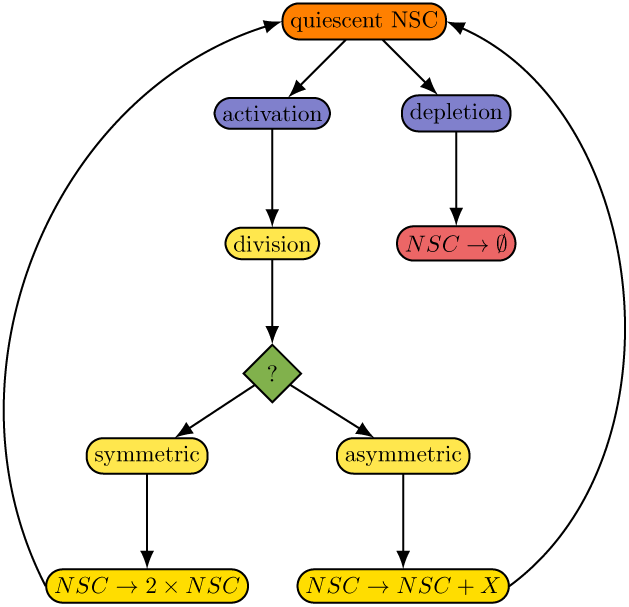
Graphical representation of the proposed model. Quiescent NSCs are either activated to enter the cell cycle and subsequently perform a symmetric or asymmetric division, or vanish from the NSC pool by a depletion event. Moreover, cycling NSCs re-enter the quiescent phase after division.

Accordingly, our model is given by the set of equations

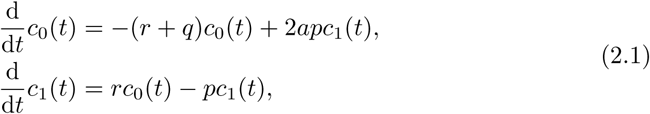

where *c*_0_ represents quiescent NSCs and *c*_1_ cycling NSCs. The parameter *p* denotes the proliferation rate, *q* is the depletion rate and *r* the activation rate (see Figure 1 for a graphical representation). Moreover, since we consider experimentally observed symmetric and asymmetric NSC divisions [8], we introduce the parameter *a* as the fraction of self-renewal, which is the probability of a daughter cell to have the same fate as a mother cell [19].

We investigate the proposed model by comparison to experimental data. For this, we measure the number of NSCs and the fraction of 5-bromo-2-deoxyuridine (BrdU) incorporating NSCs at several time points during adulthood (Figure 2). Our data agree with those reported by Encinas et al. [9], as we as well observe a decline of the NSC pool (Figure 2e) and a constant fraction of BrdU incorporating NSCs of about 1% during aging (Figure 2f). By estimating model parameters, we find that the model can be fitted to these population-level data (Figures 2e and 2f, black line).

**Figure 2:**
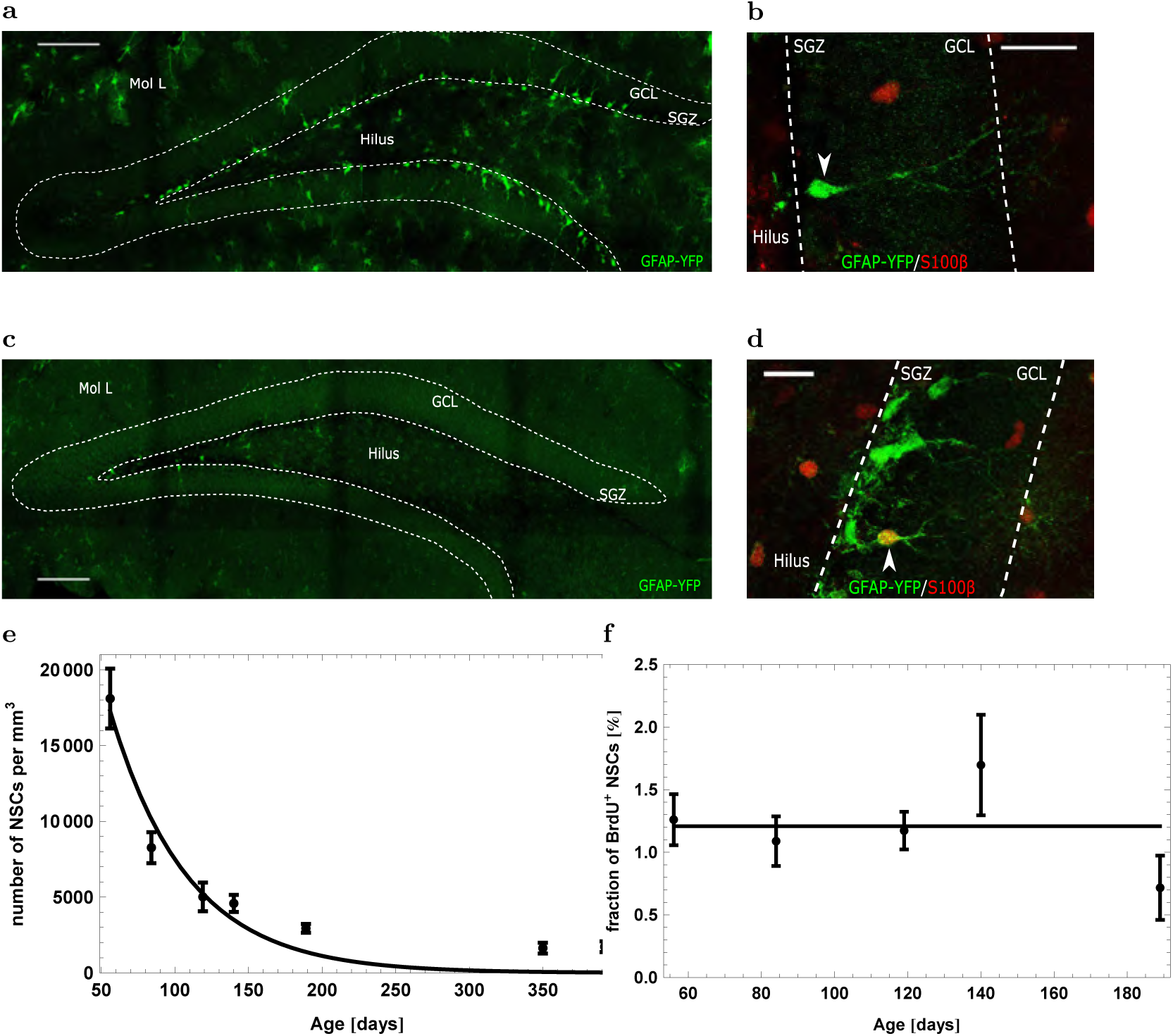
GFAP-YFP-Expressing Cells in the DG and Population-Level Dynamics of Hippocampal Neural Stem Cells. (**a**) & (**c**) GFAP-YFP-positive cells in 8-weeks-old (**a**) and 56-weeks-old (**c**) GFAP-YFP reporter mice. Scale bar 100 μm. (**b**) & (**d**)Representative confocal images of immunostaining for GFP (green) and S100*β* (red). Shown are examples of a GFAP+ /S100*β*^−^ neural stem cell (**b**) and a GFAP+/S100*β*+ astrocyte (**d**). Scale bar 20 μm. (**e**) & (**f**) Fit of the proposed model to the total number of NSCs (**e**) and the fraction of BrdU incorporating NSCs (**f**).

In contrast to population level data which account for large cell numbers and admit inferences about the collective behavior of a whole cell population, clonal data reflect single-cell level behavior by tracking the progeny of individual cells. To assess the clonal dynamics of NSCs, Bonaguidi et al. [8] labeled individual NSCs at the age of 8-12 weeks and evaluated their clonal progeny one month, two months and one year later. This led to a classification of NSC clones in three categories: *quiescent,* consisting of exactly one NSC; *activated,* including one NSC and at least one additional cell; and *depleted,* containing no NSCs.

While tissues consisting of many cells can be modeled using a deterministic approach based on averaging over the population, modeling of clonal data requires a stochastic approach taking into account cellular heterogeneity. Therefore, to fit our model to the clonal data (Figure 3), we made use of the Gillespie method Gillespie [20] to convert model (2.1) into a corresponding stochastic process.

**Figure 3:**
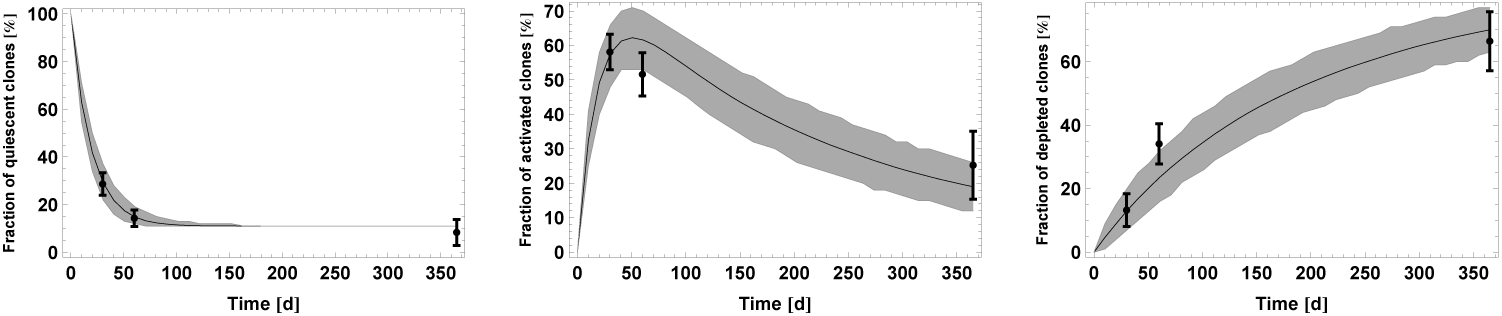
Fit of the proposed model to the clonal data of Bonaguidi et al. [8]. Results are obtained by simulating 100 NSC clones for 1000 times. Simulation data are represented as mean (solid black line) and (gray) band containing 95% of all simulated trajectories. Black error bars correspond to the clonal data.

However, the population-level and clonal dataset cannot be explained simultaneously but reflect a different dynamics, i.e. requires different parameters to reproduce the two datasets. Moreover, to simulate the clonal dynamics, it is necessary to additionally consider a population of *resilient* NSCs, cells that never become depleted or activated. The corresponding dynamics of resilient cells is outlined further on when discussing age-related effects (equation (2.3)). Assuming the existence of such subpopulation, which accounts for about 10% of all NSCs in a ten weeks old mouse, allows avoiding a rapid extinction of quiescent clones. Otherwise, the fraction of quiescent clones would approach zero and consequently the one year time point could not be matched by the model.

For a detailed description of the model quantification procedure, we refer the reader to the Materials and Methods section.

### 2.2. Aged Stem Cells Spend Longer Time in Quiescence but also Have a Higher Probability to Become Activated

The age-related decline of the NSC pool is a central characteristic of adult hippocampal neurogenesis. We asked whether the decline occurs uniformly with age or if the dynamics of NSCs change during adulthood. In case of a uniform dynamics, NSCs numbers are expected to drop exponentially, corresponding to a constant decay rate. As outlined in our previous study [21], the decline of NSC numbers saturates during aging, resulting in an underestimation of the total number of NSCs at old age (Figure 2e). This indicates that the dynamics of NSCs changes during aging. Using mathematical modeling, we evaluate different scenarios for their ability to explain this saturation.

#### 2.2.1. Decreasing Depletion

The depletion process is the integral part of the NSC decline. Without it, the number of NSCs would not decrease. One possibility to explain the saturation of NSC's decline is that the depletion rate of NSCs declines during aging, leading to a decreased fraction of depleting stem cells at old age. Accordingly, we modify model (2.1) by assuming

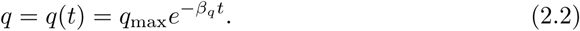

As can be seen from Figure 4a, the discussed mechanism is a very good explanation for the data.

**Figure 4:**
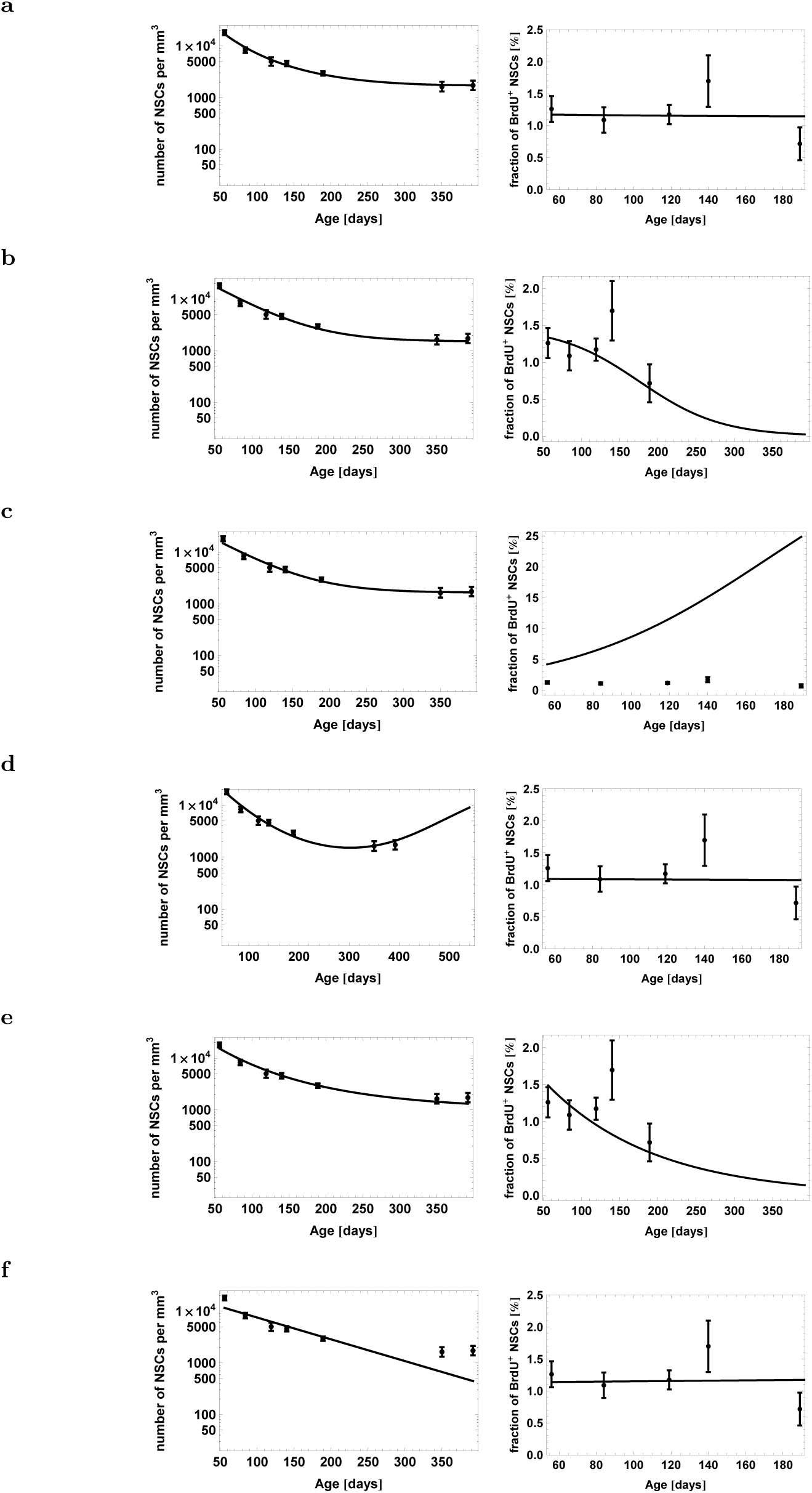
Evaluation of different scenarios to explain the saturation of the stem cell decline. Model fit to the population-level data of Figure 2, assuming (**a**) a decreasing NSC depletion rate during aging, (**b**) an additional population of resilient NSCs, (**c**) age-related lengthening of NSC's cell cycle, (**d**) that stem cells increase their self-renewal during aging, (**e**) that stem cells stay progressively longer in quiescence during aging, (**f**) that the fraction of activated stem cells increases during aging.

#### 2.2.2. Existence of Resilient NSCs

Our preceding analysis of the clonal data set of Bonaguidi et al. [8] points towards a possible second population of resilient NSCs, cells that can neither become activated nor deplete. The additional population *c*_res_ is implemented with

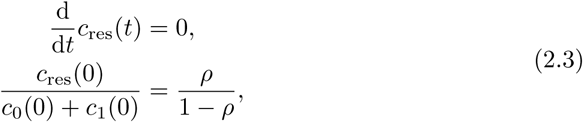
where *ρ* is the fraction of resilient NSCs on all NSCs at the start of adulthood. However, the existence of such population in combination with the decline of NSCs leads to a decrease of the fraction of cycling NSCs (
Figure 4b), contradicting the observed constant fraction of BrdU incorporating NSCs at old age [9].

#### 2.2.3. Lengthening of the Cell Cycle

If NSCs take progressively longer during aging to complete the cell cycle, the number of stem cells entering the quiescent phase declines with time, leading to a decreasing net depletion of stem cells residing in quiescence. Accordingly, we assume a decline of the proliferation rate given by

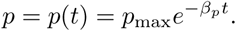

A comparison of the suggested mechanism with the data shows a lack of fit, because lengthening the cell cycle dramatically increases the fraction of cycling NSCs and alongside the fraction of BrdU incorporating NSCs (Figure 4c).

#### 2.2.4. Increasing Self-Renewal

The saturation of the NSC decline could also indicate that NSCs increase their selfrenewal to counteract the depletion. The corresponding modification takes the form

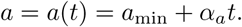

An analysis of this scenario shows that in order to explain our population level data, NSC numbers would start to increase after about 1 year of age (Figure 4d). However, this implication contradicts the fact that NSCs and other downstream compartments such as neural progenitors and immature neurons decline in numbers even at later time points than one year of age [9, 4].

#### 2.2.5. Increasing Quiescence

An increase of NSC's quiescence, corresponding to an age-related lengthening of the *G*_0_ phase, could also explain the decline pattern of NSCs. Since leaving the quiescent phase is associated with a higher tendency to deplete than to maintain or expand the pool of NSCs—otherwise it could not be explained why this pool declines—remaining in quiescence would neutralize the decline. The modification takes the form

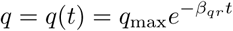

and

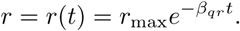

The justification for these equations is that leaving the quiescent phase *c*_0_ is driven by a joint decay process [22], which consists of activation (with rate *r*) and depletion (with rate *q*). The mean time of NSCs to sojourn in quiescence is thus given by

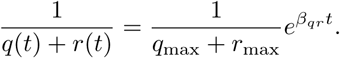

However, in order to explain the saturation of NSC's decline in this scenario, the strong increase of quiescence reduces the fraction of cycling NSCs (Figure 4e), contradicting the observation of a constant fraction of BrdU incorporating stem cells up to even 20 months of age [9].

#### 2.2.6. Increasing Activation

Another possibility to explain the saturation of the NSC decline is that the fraction of quiescent NSCs that become activated per time unit increases during aging. Since activation is associated with maintenance or expansion of the NSC pool, increasing activation could counteract the NSC decline at old age. The increasing activation scenario is implemented with

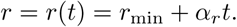

However, comparing the above mechanism to our population level data shows that an increase of activation fails to explain the saturation of NSC's decline (Figure 4f).

#### 2.2.7. Model Selection

In conclusion, the only biologically plausible scenario by visually comparing the model fit to the data is that of a decreasing depletion rate. To also evaluate each of the above model modifications from a statistical perspective, we compute for each scenario the corresponding Akaike information criterion (AIC) score. This model selection score quantifies the trade off between the complexity of a model, i.e. number of parameters, and the goodness of fit to the data [23]. Using this score, the only two considerable mechanisms besides the decreasing depletion mechanism are the scenario of increasing self-renewal and the existence of a population of resilient NSCs (see Materials and Methods). However, as outlined previously, the former scenario contradicts the observation of a decline of NSCs and downstream cell types at old age [4], while the latter contradicts the observation of a constant fraction of BrdU incorporating NSCs up to 20 months of age [9]. These considerations lead to the conclusion that among the contemplated mechanisms, the decreasing depletion scenario is the only plausible explanation for the saturation pattern of the NSC decline.

#### 2.2.8. Biological Interpretation

Only a decreased depletion rate of NSCs is able to reproduce the data without making biologically implausible predictions. A further quantification of this scenario suggests that aged stem cells stay longer in the quiescent stage (Figure 5a), but also have a higher probability of reactivation (Figure 5b). Although the two statements may seem contradictory, our model assumes that leaving quiescence can be achieved through activation or depletion. Thus, the latter assertion merely states that the probability of exiting the quiescent stage through activation versus depletion increases during aging.

**Figure 5:**
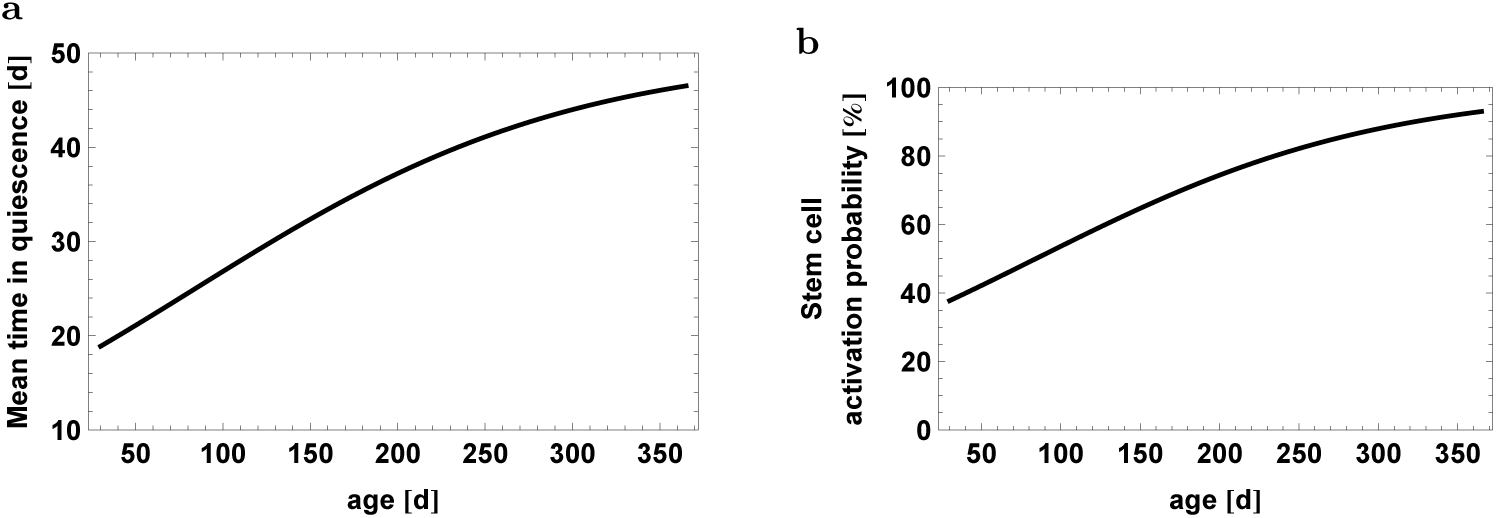
Predicted age-related changes of (**a**) the average time a stem cell stays in the quiescent phase and (**b**) the probability a stem cell becomes activated rather than depleted from the quiescent phase.

##### Stem Cell Depletion is Driven by Astrocytic Transformation and Apoptosis

In order to explain the decline of NSC numbers during aging, Encinas et al. [9] suggested that NSCs deplete by transforming into astrocytes. To test this theory, we counted the total number of astrocytes during aging to see whether astrocytes accumulate in the DG due to stem cell depletion. We also checked whether newborn astrocytes migrate away from the granule cell layer and escape counting. To this end, we used sparse labeling of Nestin-YFP reporter mice that exclusively marked NSCs within the subgranular zone. We did not observe any YFP positive cells outside the dentate gyrus 90 days after the last tamoxifen injection. Since we find an increasing progression of the astrocyte count (Figure 6), we use our model to test whether the increase in astrocyte numbers corresponds to the observed stem cell decline. For this, we take model (2.1) together with our best explanation of the saturation of the NSC decline (2.2) and add a new compartment c_2_ of astrocytes satisfying 
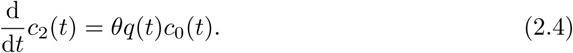

**Figure 6:**
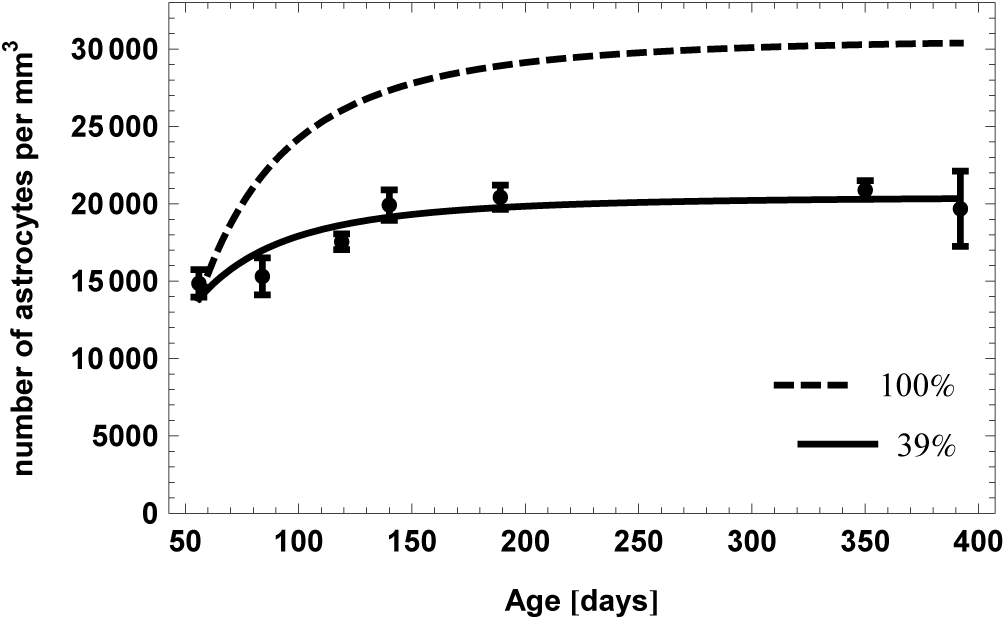
Age-related accumulation of astrocyte numbers. Dashed line represents expected number of astrocytes, assuming that 100% of the NSC decline is caused by astrocytic transformation. Solid line results from fitting the fraction of transformation events to the data, indicating that ~ 40% of the NSC decline is caused by astrocytic transformation.

Here, *θ* ∈ [0,1] is the fraction of NSC depletion events where depletion occurs via astrocytic transformation. An estimation of this parameter yields *θ* = 0.391. Thus, modeling reveals that astrocytic transformation only partially contributes to about 40% of the stem cell decline. If every NSC (100%) depletes through transformation into astrocytes, the final astrocytic yield would be much higher than what is observed (Figure 6). The remaining 60% of the NSC decline are likely a result of apoptosis and further modeling laid out in the appendix shows that NSC apoptosis is almost non-detectable due to rapid phagocytosis by microglia [11].

It is also possible that the accumulation of astrocyte numbers can be explained with a higher transformation probability *θ*, if additionally astrocytes are allowed to undergo apoptosis, thus counteracting the total astrocytic yield. We also analyzed this scenario in the appendix and find that apoptosis cannot account for a higher transformation probability.

### 2.3. Upon Dkk1 Deletion, Stem Cells Spend Longer Time in Quiescence but Are More Likely to become Re-Activated

We apply our quantified model of NSC dynamics to the study of Seib et al. [6], in which the Wnt antagonist Dickkopf-1 (Dkk1) was deleted in NSCs. The deletion led to a larger NSC pool that even counteracted the age-related decline [6]. In view of published studies [24, 25], we previously hypothesized that NSC numbers expand by increasing the fraction of self-renewal in response to increased Wnt activity. Now, we used the developed model to identify the most likely change in cellular dynamics that explains Wnt-induced expansion of the pool of NSCs.

First, we extend our stem cell model (2.1) including aging effects (2.2) and astrocytic transformation (2.4) by adding all cell types necessary to simulate hippocampal neurogenesis. In particular, we assume that (i) NSCs generate progenitors and astrocytes via asymmetric divisions [8], (ii) progenitors perform a sequence of symmetric divisions followed by differentiation into neuroblasts [9] and (iii) neuroblasts undergo apoptosis as well as neuronal differentiation [11]. Thus, the full model of wild type neurogenesis is given by the set of equations

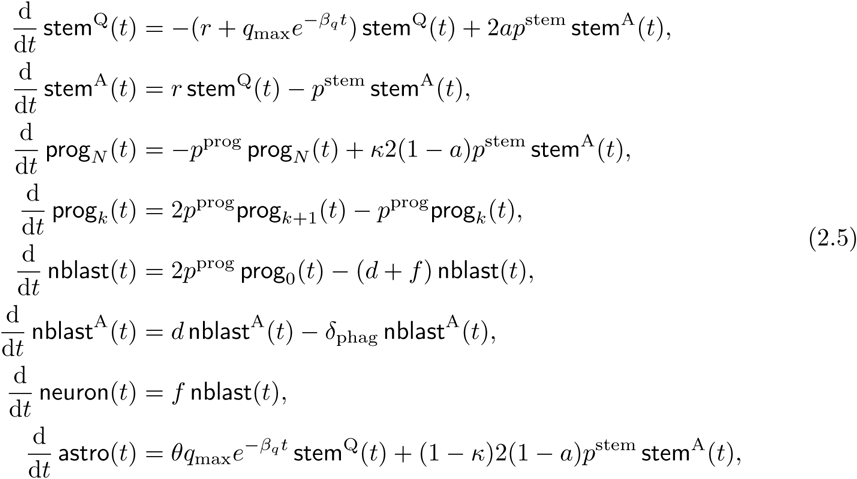

where stem^Q^ denotes quiescent stem cells, stem^A^ active (cycling) stem cells, prog_*i*_ progenitors with *i* remaining divisions (1 ≤ *i* ≤ *N*), nblast^A^ neuroblasts, nblast^A^ apoptotic neuroblasts, neuron mature neurons and astro astrocytes.

Next, we use the full model to simulate the protocol of the Dkk1 study (Figure 7a). In the experiment, adult mice were injected with Tamoxifen (TAM) to induce deletion of Dkk1. Following a 5 week period, BrdU was administered and chased for 24 hours and 4 weeks. The number of BrdU labeled cells among the different populations following Dkk1 deletion was quantified and compared to numbers in wildtype animals.

**Figure 7:**
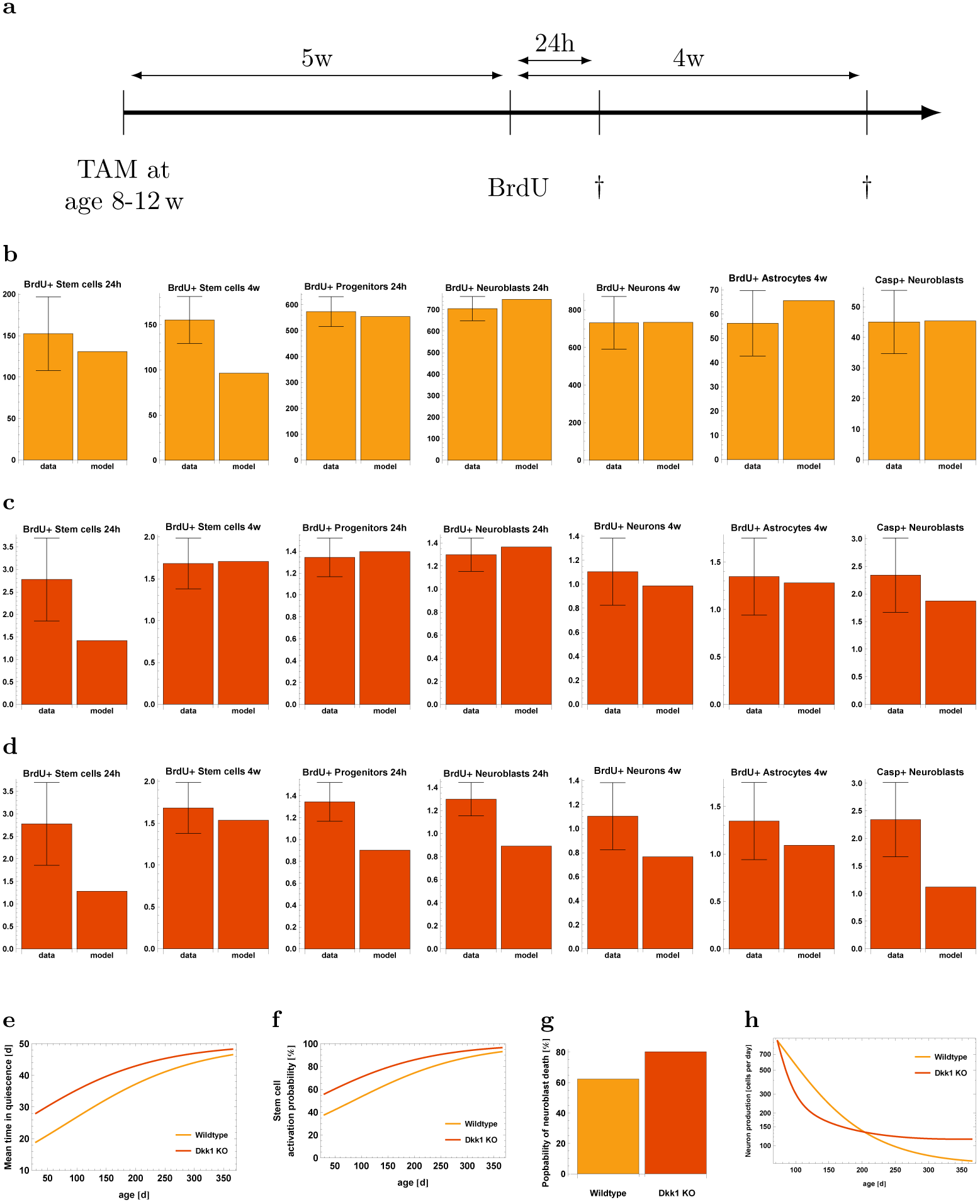
Evaluation of the Dkk1 Knockout Study [6]. **(a)** Graphical representation of the experimental protocol **(b)** Comparison of the wildtype part of the data and the mathematical model **(c)** Observed KO effects versus model, assuming decreased NSCs depletion rate. KO effects are defined as ratio of the KO data to the corresponding WT data, i.e. KO effect = (KO data) / (WT data) **(d)** Observed KO effects versus the model, assuming increased NSCs self-renewal **(e)** Mean time of a stem cell to remain in the quiescent stage. In the case of Dkk1 deletion, this time is increased. **(f)** Age-dependent activation probability of stem cells. In the KO case, NSCs have a higher activation probability. **(g)** Probability of a neuroblast to die rather than maturing into a neuron. In the case of Dkk1 knockout, a higher fraction of neuroblasts dies. **(h)** Production rate of new neurons. Dkk1 deletion alters the number of newborn neurons and results in higher neuron production at old age.

In order to explain the effects of Dkk1 deletion, we first reproduced the wild-type (WT) part of the data using our model. As can be seen from Figure 7b, the model displays a very good fit to the WT data. Interestingly, a prediction of this fit is that 38% of the neuroblasts mature into neurons and about 62% undergo apoptosis. This finding is consistent with experimental observations stating that the major part of immature neurogenic progenitors undergoes apoptosis rather than differentiating into neurons [11], thus further supporting our modeling approach.

We then focus on explaining the effects of Dkk1 knockout (KO). To model the effects of Dkk1 deletion, we introduced additional parameters into the wild type model (2.5). Since Dkk1 was selectively deleted in NSCs, we assume that knockout effects were the result of changes in the value of stem cell parameters. We thus consider the parameters *a*, *p*^stem^, *q*_max_ and *r* as possible candidates for being changed upon Dkk1 deletion. In addition, the data indicates an increased death rate *d* of neuroblasts after the knockout. For each candidate parameter 
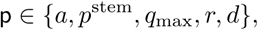
we introduce a change Δ_p_ such that the corresponding wild-type and knockout parameters are related via

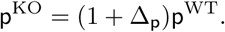

Hence, Δ_p_ denotes the relative change of the WT parameter p. Moreover, since the model fit to the WT part of the data is not perfect, instead of considering the KO data we consider KO *effects,* defined as the ratio of the KO data to the corresponding WT data, i.e. KO effect = (KO data)/(WT data).

We then apply statistical measures to analyze different combinations of the candidate parameters for their plausibility of being altered simultaneously by Dkk1 deletion using AIC scores. We find that the most plausible explanation of the observed changes is a decreased depletion rate *q*_max_ in combination with increased neuroblast apoptosis d (Figure 7c). In contrast, statistical analysis shows that alternative scenarios such as increased NSC self-renewal (Figure 7d) can be discarded as explanation of the KO data (see Materials and Methods). To quantify the above mentioned explanation of decreased NSC depletion rate *q*_max_ and increased neuroblast apoptosis rate *d*, we calculate the average time a stem cell remains quiescent (Figure 7e) and the probability that leaving quiescence is driven by activation, i.e. cell cycle entry, rather than depletion (Figure 7f). In young individuals, stem cells remain about ten days longer quiescent and have a 20 percentage points increased activation probability in the case of Dkk1 deletion, while those differences diminish during aging. Moreover, Dkk1 knockout increases the probability that neuroblasts undergo apoptosis rather than neuronal differentiation from about 60% to about 80% (Figure 7g). Interestingly, Dkk1 deletion impacts the production rate of new neurons (Figure 7h): In young individuals, neuron production is decreased due to increased NSC quiescence. The latter leads to increased NSC numbers in old individuals, that in turn results in increased neuron production.

## 3 Discussion

To obtain insights into the control mechanisms of stem cell dynamics and age-related effects during hippocampal neurogenesis, we have applied an interdisciplinary approach based on mathematical modeling and experimental data. This approach allows investigating normal and perturbed neurogenesis by quantifying cell dynamics that cannot be measured experimentally.

Our stem cell model (2.1) was motivated by the experimental observations of Bonaguidi et al. [8] and subsequently extended combining data on NSCs cell cycle properties [12], astrocytic transformation (newly generated), progenitor dynamics [9] as well as neuroblast dynamics and phagocytosis [11]. Thus, our modeling framework integrates data describing neurogenesis on different levels in a coherent manner.

Although the original stem cell model (2.1) can reproduce the available population-level and clonal data, it should be noted that both datasets reflect different dynamics corresponding to different cell parameters. There might be several reasons for such discrepancy. Different morphological and genetic features used to define NSCs may correspond to distinct sub-populations of stem cells or conversely, to slightly different differentiation stages within a continuum between a stem cell and a neural progenitor. In addition, the clonal data account for relatively few cells collected from multiple individuals. Thus, averaged clonal data as in the case of Bonaguidi et al. [8] could be biased due to biological variability among animals or certain cell fate choices stochastically dominating due to the paucity of analyzed clones.

We next addressed for possible mechanisms that can explain the saturation pattern of the age-related decline in NSC numbers—an observation that confirms previous data by Encinas et al. [9]. After evaluating several stem-cell features, the only explanation left is that stem cells spend a progressively longer time in quiescence during aging. At the same time, a greater fraction of aged NSCs becomes re-activated from the stem cell pool rather than being depleted. Our reasoning is coherent from a modeling perspective, since we have ruled out several alternative scenarios.

Another question related to age-related changes during hippocampal neurogenesis is that of the mechanism leading to the decline of NSC numbers. Here, we were able to partially confirm the astrocytic transformation hypothesis of Encinas et al. [9] by observing an age-related accumulation of astrocyte numbers. Mathematical modeling revealed that these transformation events account for about 40% of the NSC decline. Interestingly, our calculations showed that even if the remaining 60% of the decline is caused by NSC apoptosis, detection of apoptotic cells is not possible due to rapid phagocytosis [11]. Similarly high estimates of apoptosis have been obtained in a modeling study of neurogenesis based on a stochastic branching process approach [26].

One of the main aims of mathematical modeling is to develop model-based predictions which can be tested experimentally. Such model validation is an important step within the scientific method [27]. We thus applied our model to the study of Seib et al. [6], in which the Wnt inhibitor Dkk1 was deleted. The very good fit to the wild-type data, which results from accurate simulation of the experimental protocol, confirms that our model accounts for key aspects of hippocampal neurogenesis. Moreover, the fitting procedure has led to the prediction that the major part of neuroblasts undergoes apoptosis instead of neuronal differentiation, a prediction which has already been validated [11]. Moreover, we have analyzed multiple scenarios to reveal the effects of Dkk1 deletion. Originally, Seib et al. [6] hypothesized that Dkk1 knockout increases the self-renewal of NSCs. This reasoning was based on previous studies showing that Wnt signaling induces self-renewal of radial glia progenitors in the embryonic brain [25] and that Wnt ligands were reported to increase the proliferation and self-renewal of NSCs [28, 29, 30, 24]. However, modeling shows that the only considerable scenario is that of an increased NSC quiescence in conjunction with an increased probability to exit the quiescent phase through activation rather than depletion. In contrast, increased self-renewal of NSCs can not explain the effects upon Dkk1 knockout. The reason for this is that increasing the self-renewal of NSCs would increase the fraction of quiescent NSCs on all NSCs, since our model assumes NSCs return to quiescence after division [8], and alongside a decreased fraction of BrdU positive downstream cell compartments would be observed. Taken together, our results indicate that regulating the rate of activation vs. depletion into astrocytes seems to represent a central hub to fine-tune neurogenesis.

Our study shows that mathematical modeling is a powerful tool to investigate complex cell systems such as the neurogenic niche of the hippocampus. It is possible to describe the dynamics of stem cell systems using both deterministic [31, 32, 33] and stochastic models [34, 35, 16]. While the former usually have the form of linear or nonlinear differential equations, the latter are of at least two possible kinds. One is the system of master equations (Chapman-Kolmogorov forward equations if the model is Markovian), which allow computing probability distributions of the state variables as a function of time. If it is desirable to obtain longitudinal information, i.e. information concerning the possible time trajectories of the process, the most common practical solution is simulation. A fusion of deterministic and stochastic simulation approaches, as conducted in this study by employing the Gillespie algorithm [20], is very effective to integrate population-level and single cell-level data. The initial deterministic stem cell model consists of linear differential equations, featuring constant rates. Such linear models have been successfully applied before to processes close to homeostasis [36] or uncontrolled growth in cancer [37, 33]. In our case, using a linear model allowed identifying time-dependent parameters necessary to explain age-related changes during neurogenesis. This, in turn, suggests which parameters are subject to a nonlinear regulation.

## 4 Materials and Methods

### 4.1. Animals

GFAP-CreER^T2^-YFP and Nestin-CreER^T2^-YFP reporter mice were housed in the animal facilities of the German Cancer Research Center (DKFZ) at a 12 h dark/light cycle with free access to food and water. All animal experiments were performed in accordance with the institutional guidelines of the DKFZ and were approved by the Regierungspräsidium Karlsruhe, Germany.

### 4.2. Tamoxifen and BrdU Administration

GFAP-CreER^T2^-YFP mice received an intraperitoneal (i.p.) Tamoxifen (Sigma; T5648) injection (40mg/kg) twice a day (morning and evening), for 5 consecutive days. Two days after Tamoxifen treatment, a single shot BrdU (Sigma; B5002; 150mg/kg) was i.p. injected and mice were sacrificed two hours later.

### 4.3. Long-Term Tracing of NSC-Derived Astrocytes

Nestin-CreER^T2^-YFP reporter mice received Tamoxifen injections as described above and were sacrificed 90 days after the last injection. Thereafter the presence of YFP labeled cells outside the hippocampal region was examined. We did not find any YFP positive cells outside the DG granule cell layer.

### 4.4. Tissue Preparation and Staining

Animals were perfused with 1×HBSS (Gibco; 14170-088) and 4% PFA (Carl Roth; P087.1). Subsequently the brain was postfixed overnight in 4% PFA and tissue was cut in 50μm thick coronal brain slices using a Leica VT 1200S vibratome.

### 4.5. Immunohistochemistry

For each animal, six 50 μm thick coronal brain sections (250 μm distance between sections) were washed four times, 10 minutes in TBS at room temperature, blocked for one hour in TBS containing 3% horse serum (Millipore; S9135) and 0.3% Triton X-100 (Sigma; 9002-93-1) at room temperature, followed by an overnight staining at 4°C in TBS containing 3% horse serum and 0.3% Triton X-100 with the following primary antibodies: rat anti BrdU (Abcam; ab6326), chicken anti GFP (Aves; GFP-1020), mouse anti S100 *β* (Abcam; ab66028) and rabbit anti Tbr2 (Abcam; ab23345). The next day, brain sections were washed four times, 10 minutes in TBS at room temperature, blocked for 30 minutes TBS containing 3% horse serum and 0.3% Triton X-100 at room temperature and stained for two hours in TBS containing 3% horse serum and 0.3% Triton X-100 at room temperature with the following secondary antibodies: donkey anti rat 405 (Abcam; ab175670), donkey anti chicken 488 (Jackson ImmunoResearch; 703-545-155), donkey anti mouse 549 (Jackson ImmunoResearch; 715-507-003) and donkey anti rabbit 647 (Jackson ImmunoResearch; 711-605-152). Afterwards sections were washed four times, 10 minutes in TBS at room temperature and mounted on glass slides.

### 4.6. Imaging and Quantification

For each dentate gyrus, confocal z-stacks (2 μm distance between images) were acquired with a Leica TCS-SP5 confocal microscope with a 20x oil immersion objective and a resolution of 1024 × 1024 at 100 Hz. Obtained images were analyzed using the ImageJ Cell Counter Plugin to manually count and mark single cells of different cell types. Different cell populations were defined using the following marker combinations: Neural stem cells (YFP+/S100*β*^−^/Tbr2^−^), astrocytes (S100*β*+/Tbr2^−^), and cycling cells that gained the marker BrdU by retaining their markers described above. Determined cell numbers were quantified as number of cells per cubic millimeter in the DG granule cell layer. The DG volume was calculated by the area of the DG on the central image of the z-stack (measured with ImageJ), multiplied with the z-stack size.

### 4.7. Mathematical Modeling

#### 4.7.1. Quantification

The stem cell model (2.1) contains four parameters *p, q, r* and a. The value of the proliferation rate *ρ* can be inferred from the literature [12], since the length of the cell cycle of NSCs was measured as

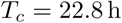

and we can interpret this value as the doubling-time of an exponential growth process, leading to

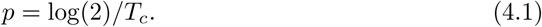

For the fraction of self-renewal *a* we assume that 2*a −* 1, the corresponding probability of a division being symmetric [21], is 5%. We justify this choice of a with the suggestion of Bonaguidi et al. [8] that the fraction of symmetric NSC divisions on all divisions is relatively small. A different value of *a* does not affect the drawn conclusions.

The remaining two free parameters *q* and *r* are estimated from the population level data of the total number of stem cells (Figures 2e) and the fraction of BrdU incorporating stem cells (Figure 2f). To fit the BrdU incorporation data, we use that the fraction of BrdU incorporating NSCs is the product of the fraction of cycling NSCs on all NSCs times the relative length of the S-phase in the cell cycle, *T_s_/T_c_,* with *T_s_* = 9.7h [12]. The reason is that the thymidine analog BrdU can only be incorporated in cells which are in the stage of DNA synthesis.

In the case of estimating *q* and *r* from the clonal data (Figure 3), the model was converted a into corresponding stochastic processes using the Gillespie method [20]. Afterwards, 100 NSC clones were simulated for 1000 times and the resulting mean fraction of quiescent, activated and depleted clones was fitted to the clonal data.

In general, model fitting was performed using the `NonlinearModelFit` procedure of *Mathematica* 9 to numerically minimize the weighted sum of squared residuals. Weights were chosen as inverse squares of the data points standard error of the mean according to the *Mathematica* documentation for fitting data with measurement errors. Estimated parameters for the stem cell model are summarized in the Table 1.

**Table 1:** Estimated parameters of model (2.1). Data type indicates whether the model was fitted to the population-level data (Figures 2e and 2f) or the clonal data (Figure 3).

| Data Type | Parameters |
| --- | --- |
| Population-level | $q = 0.021 \text{ d}^{-1}$<br>$r = 0.021 \text{ d}^{-1}$<br>$\rho = 0$ |
| Clonal | $q = 0.0057 \text{ d}^{-1}$<br>$r = 0.047 \text{ d}^{-1}$<br>$\rho = 0.11$ |

The full neurogenesis model (2.5) contains 12 parameters, of which 9 are already estimated or taken from the literature (Table 2). The parameters *κ*, *d* and *f* are not known prior to the Dkk1 experiment and need to be inferred from fitting the full model to the WT part of the data.

**Table 2:**
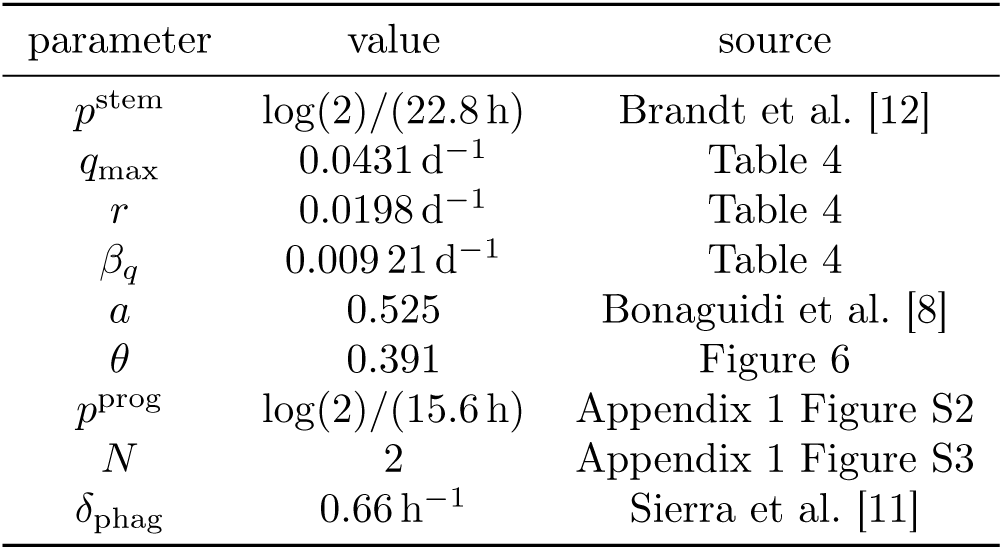
Parameter values of system (2.5), which is used to model the Dkk1 experiment of Seib et al. [6].

Estimating the unknown parameters *κ*, *d* and *f* leads to the values displayed in Table 3. As previously stated, the resulting fraction of surviving neuroblasts,

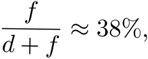

is consistent with experimental findings [11].

**Table 3:**
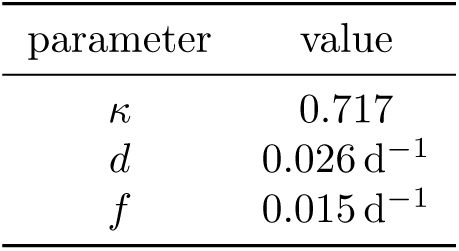
Estimated parameters of the full neurogenesis model (2.5) as a result of fitting the WT part of the Dkk1 data [6].

#### 4.7.2. Model Selection

To assess the plausibility of the different scenarios to explain the saturation of the NSC decline, we make use of model selection theory. For each scenario, we compute its corresponding AIC value and compare the resulting Akaike weights Δ (Table 4), which penalize overly complex models. The recommendation is that the level of empirical support of a certain model is substantial if 0 ≤ Δ ≤ 2, considerably less, if 4 ≤ Δ ≤ 7 and essentially none, if Δ > 10 holds [23]. Thus, the only two considerable mechanisms besides the decreasing depletion mechanism are the scenario of increasing self-renewal and the existence of a population of resilient NSCs (Table 4). However, as outlined previously, the former scenario contradicts the observation of a decline of NSCs and downstream cell types at old age [4], while the latter contradicts the observation of a constant fraction of BrdU incorporating NSCs up to 20 months of age [9].

**Table 4:** Parameters estimated during the analysis of different mechanisms to explain the saturation of the NSC decline. AICc is the small sample size corrected Akaike information criterion and ΔAICc the corresponding Akaike weight, i.e. the difference to the smallest AICc value.

| Mechanism | Parameters | $R^2$ | AICc | $\Delta\text{AICc}$ |
| --- | --- | --- | --- | --- |
| Decreasing depletion | $q_{\max} = 0.043 \text{ d}^{-1}$<br>$r = 0.020 \text{ d}^{-1}$<br>$\beta_q = 0.0092 \text{ d}^{-1}$ | 0.915 | 242.26 | 0 |
| Resilient population | $q = 0.021 \text{ d}^{-1}$<br>$r = 0.026 \text{ d}^{-1}$<br>$\rho = 0.035$ | 0.909 | 245.96 | 3.7 |
| Cell cycle lengthening | $q = 0.020 \text{ d}^{-1}$<br>$r = 0.027 \text{ d}^{-1}$<br>$\beta_p = 0.019 \text{ d}^{-1}$ | 0.909 | 246.26 | 4 |
| Increasing self-renewal | $q = 0.026 \text{ d}^{-1}$<br>$r = 0.019 \text{ d}^{-1}$<br>$\alpha_a = 0.0022 \text{ d}^{-1}$ | 0.907 | 247.22 | 5 |
| Increasing quiescence | $q_{\max} = 0.031 \text{ d}^{-1}$<br>$r_{\max} = 0.038 \text{ d}^{-1}$<br>$\beta_{qr} = 0.0071 \text{ d}^{-1}$ | 0.903 | 249.62 | 7.4 |
| Increasing activation | $q = 0.011 \text{ d}^{-1}$<br>$r_{\min} = 0.019 \text{ d}^{-1}$<br>$\alpha_r = 4.5 \times 10^{-6} \text{ d}^{-1}$ | 0.865 | 267.59 | 25.3 |

To find the best explanation for the effects upon Dkk1 KO, we employ a nested approach by considering simple, i.e. few parameters involving, explanations as well as more complex scenarios (Table 5). At first, we assume that only one of the stem cell parameters *a*, *p*^stem^, *q*_max_ or *r* and in addition the neuroblast death rate *d* changes. The best fit to the data is achieved by a decrease of the depletion rate *q*_max_ or an increase of the activation rate *r*. In contrast, the scenario of increased self-renewal *a* displays a considerably worse fit. The decreased depletion and increased activation scenario both lead to a shift of the balance between NSC activation and depletion towards a higher fraction activation events. We thus also consider a scenario in which both parameters, *q*_max_ and *r* change simultaneously and find that it improves the fit. It is also possible that all NSC parameters change, resulting in an even better fit.

**Table 5:** Estimated changes of WT parameters to account for the Dkk1 KO effects. ΔAICc denotes the Akaike weight of the respective model corresponding to the small-sample-size corrected Akaike information criterion (AICc).

| $\Delta_a$ | $\Delta_{p^{\text{stem}}}$ | $\Delta_{q_{\text{max}}}$ | $\Delta_r$ | $\Delta_d$ | $R^2$ | AICc | $\Delta\text{AICc}$ |
| --- | --- | --- | --- | --- | --- | --- | --- |
| .20 | — | — | — | .69 | .860 | 280.0 | 28.8 |
| — | -.20 | — | — | .75 | .819 | 292.9 | 41.7 |
| — | — | -.52 | — | 1.46 | .922 | 251.2 | 0 |
| — | — | — | .45 | 1.50 | .915 | 255.0 | 3.8 |
| — | — | -.41 | .092 | 1.49 | .922 | 253.2 | 2 |
| .055 | -.18 | -.09 | .31 | 1.39 | .924 | 257.1 | 6 |

Although the fit improves with an increasing number of parameters, we again compute AIC values to assess if the increased goodness of fit justifies the additional complexity of the model. Applying the previously mentioned recommendation of 0 ≤ Δ ≤ 2, scenarios leading to the discussed shift of the balance between NSC activation and depletion towards a higher fraction activation events should be considered first (Table 5). Moreover, the originally suggested increased self-renewal [6] of NSCs can be disregarded to explain the effects of Dkk1 deletion.

## Acknowledgments

The authors gratefully acknowledge the Collaborative Research Center, SFB 873 “Maintenance and Differentiation of Stem Cells in Development and Disease”. AM-C and FZ were also supported by the European Science Foundation (Starting Grant No. 210680) and the Emmy Noether Programme of the German Research Council (DFG).

## Supplementary Information

### A. Astrocytic Transformation

We want to give an explanation for the claim that NSC apoptosis is almost nondetectable in the case of 1 − *θ =* 60.9% of the NSC decline resulting from apoptosis. Consider the stem cell model (2.1) including aging effects (2.2). To model the dynamics of NSC apoptosis, we add a new compartment *c*_A_ accounting for *apoptotic,* i.e. biologically dead but physically present stem cells. The dynamics of NSCs at a certain age *τ* is thus given by

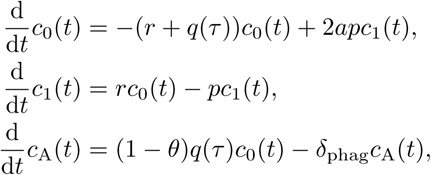

where *δ*_phag_ is the rate at which apoptotic cells are cleared via phagocytosis within, on average, *1/δ_phag_* = 1.5h [11]. At the age of *τ* = 2 months, there are about *n_0_* = 10000 NSCs in the entire dentate gyrus [9] and the number of apoptotic NSCs can be calculated by that number times the steady state fraction of apoptotic NSCs on all NSCs. Accordingly, there are

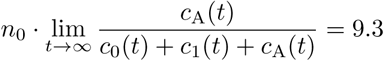

apoptotic stem cells in the entire dentate gyrus. Considering the usual sampling fraction of one sixth of the DG, there only remain about two apoptotic stem cells to be detected.

To analyze the scenario that the accumulation of astrocytes could be explained with a higher transformation rate *θ* if in addition astrocytes are allowed under go apoptosis, we consider the modified dynamics

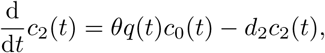

where *d*_2_ is the death rate of astrocytes. Estimating *θ* and *d*_2_ simultaneously yields *θ* = 0.392 and *d*_2_ = 2.6 × 10^−6^ *d*^−1^, showing that there is no justification for such scenario. In addition, assuming *θ* = 1 and only estimating *d*_2_ results in an AICc score of 178.9, which compared to the AICc of 144.7 for the no-apoptosis model further indicates that there is no support for this scenario from a model selection viewpoint.

### B. Dynamics of Progenitor Cells

To model the dynamics of progenitor cells, we again make use of the study of Encinas et al. [9]. We consider the data set in which the authors label dividing cells with

**Figure S1:**
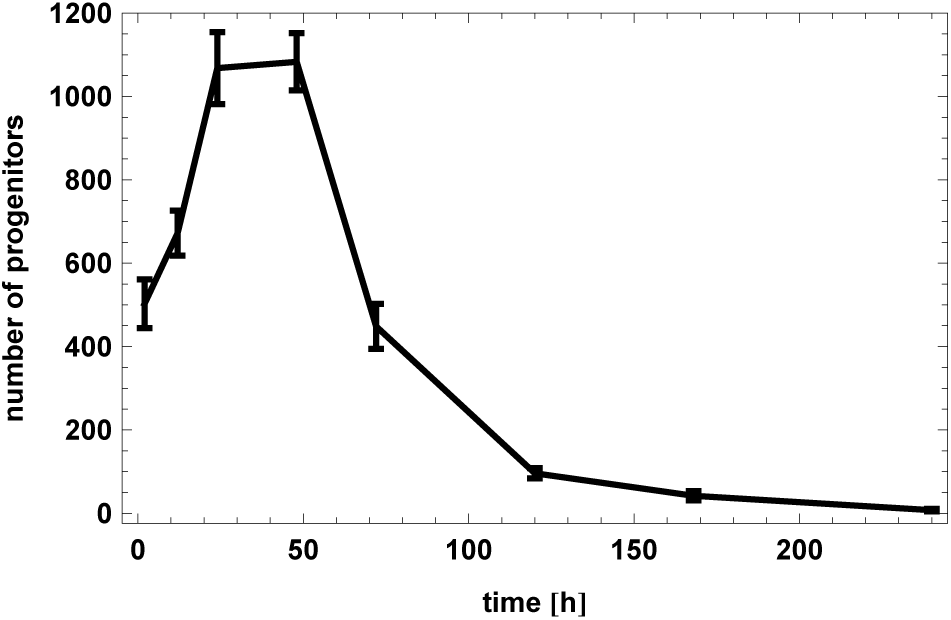
Time course of the dynamics of neural progenitor cells. Two months old mice were injected with BrdU and sacrificed at several time points after injection. Depicted is the number of BrdU positive progenitor cells. Data is reproduced from the publication of Encinas et al. [9].

BrdU and track the number of BrdU labeled progenitors (Appendix 1 Figure S1). Because of the rapid increase and subsequent decrease of labeled cells, they concluded that progenitors perform a series of symmetric self-renewing divisions, followed by subsequent transformation into neuroblasts.

We model the dynamics of progenitors with the equations

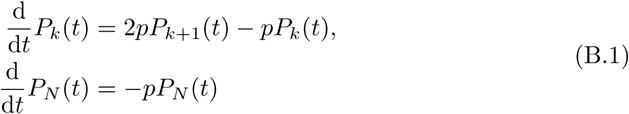

for *k =* 0*,…,N −* 1. Here, *P_i_* is the number of progenitor cells with *i* remaining divisions and *p* > 0 is the proliferation rate. Moreover, we assume that at the start of the experiment, all progenitors have *N* remaining divisions, i.e. *P_N_*(0) = *n* for some *n >* 0 and *P_k_* (0) = 0 for *k ≠ N*.

For quantifying *p*, we consider the corresponding cell cycle length *t_c_*, which is linked to *p* via (4.1). The cell cycle length of progenitor cells has been measured in different studies, however with contrasting results ranging from 12 − 14 h to about 22 h [38, 39]. We thus employ an unbiased approach for quantification by assuming different cell cycle lengths *t_c_* and compute the *R*^2^ of the fit dependent on *N*, the maximum number of progenitor divisions (Appendix 1 Figure S2).

The best fit can be obtained for *N* =2, 3 or 4, but only *N* = 2 allows for a cell cycle length in the range of what is experimentally observed with the maximum *R*^2^ at *t_c_* = 14.4 h. To achieve a better compromise between our model assumption and the measured cell cycle lengths, we relax the condition of an optimal *R*^2^. A visual assessment of the fit shows that *R*^2^ = 0.95 provides a reasonable fit to the data. For *N* =2, the maximal *t_c_* for which *R*^2^ = 0.95 can be achieved is *t_c_* = 15.6 h (Appendix 1 Figure S3), which we assume for our subsequent analysis.

**Figure S2:**
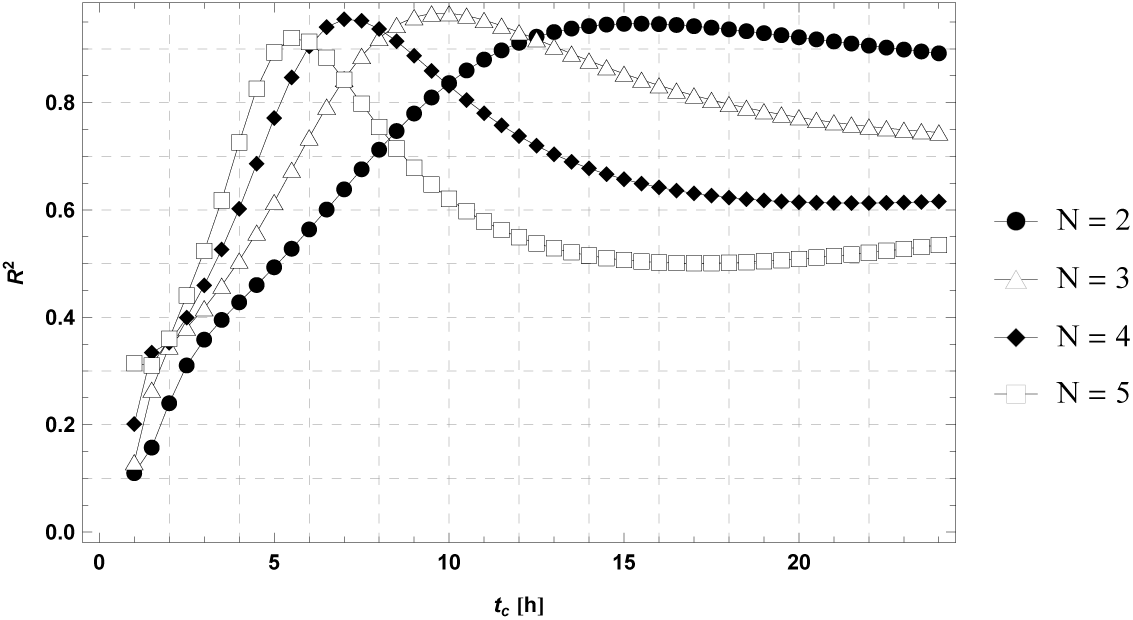
Goodness of fit of the progenitor model (B.1) to the data displayed in Appendix 1 Figure S1. The *R*^2^ is calculated solely from the first five time points of the data, which capture the rapid rise and fall of progenitor numbers.

**Figure S3:**
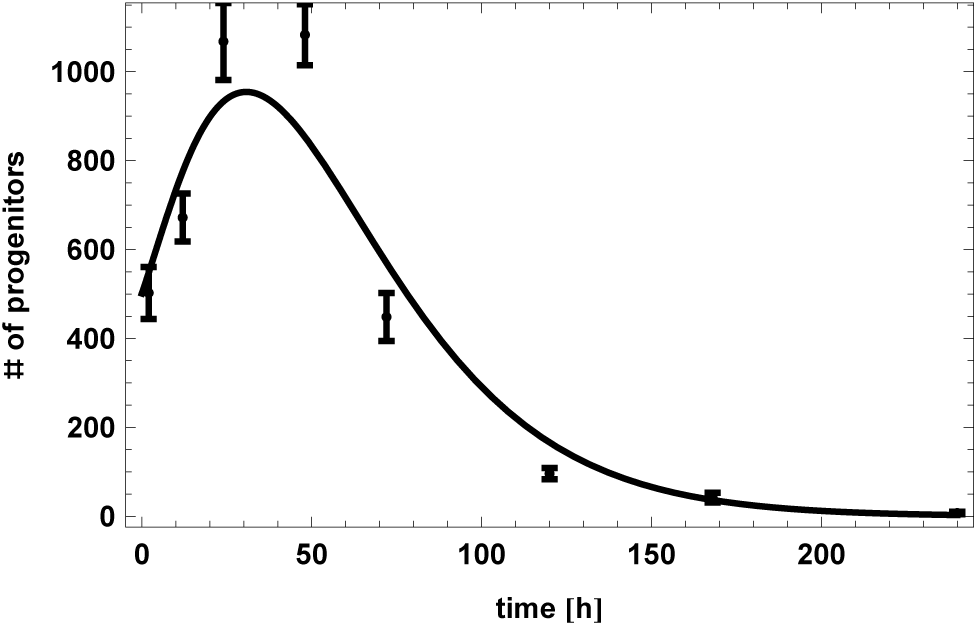
Fit of the progenitor model (B.1) to the data of Appendix 1 Figure S1, assuming *N = 2* progenitor divisions and a cell cycle length of *t_c_* = 15.6h.

## References

[1] Guo-Li Ming and Hongjun Song. “Adult neurogenesis in the mammalian brain: significant answers and significant questions.” In: Neuron 70 (2011), pp. 687–702.

[2] N. M. Ben Abdallah et al. “Early age-related changes in adult hippocampal neurogenesis in C57 mice”. In: Neurobiol. Aging 31 (2010), pp. 151–161.

[3] S. Jinno. “Decline in adult neurogenesis during aging follows a topographic pattern in the mouse hippocampus”. In: J. Comp. Neurol. 519 (2011), pp. 451–466.

[4] Josephine Walter et al. “Age-related effects on hippocampal precursor cell subpopulations and neurogenesis”. In: Neurobiology of aging 32.10 (2011), pp. 19061914.

[5] S. Furutachi et al. “p57 controls adult neural stem cell quiescence and modulates the pace of lifelong neurogenesis”. In: EMBO J. 32 (Apr. 2013), pp. 970–981.

[6] D. R. Seib et al. “Loss of dickkopf-1 restores neurogenesis in old age and counteracts cognitive decline”. In: Cell Stem Cell 12 (2013), pp. 204–214.

[7] Kieran M Jones et al. “CHD7 Maintains Neural Stem Cell Quiescence and Prevents Premature Stem Cell Depletion in the Adult Hippocampus”. In: Stem Cells 33.1 (2015), pp. 196–210.

[8] Michael A. Bonaguidi et al. “In Vivo Clonal Analysis Reveals Self-Renewing and Multipotent Adult Neural Stem Cell Characteristics”. In: Cell 145 (2011), pp. 1142–1155.

[9] Juan M. Encinas et al. “Division-Coupled Astrocytic Differentiation and Age-Related Depletion of Neural Stem Cells in the Adult Hippocampus”. In: Cell Stem Cell 8 (2011), pp. 566–579.

[10] Chitra D Mandyam, Gwyndolen C Harburg, and Amelia J Eisch. “Determination of key aspects of precursor cell proliferation, cell cycle length and kinetics in the adult mouse subgranular zone”. In: Neuroscience 146.1 (2007), pp. 108–122.

[11] A. Sierra et al. “Microglia shape adult hippocampal neurogenesis through apoptosis-coupled phagocytosis”. In: Cell Stem Cell 7 (2010), pp. 483–495.

[12] M. D. Brandt, M. Hubner, and A. Storch. “Brief report: Adult hippocampal precursor cells shorten S-phase and total cell cycle length during neuronal differentiation”. In: Stem Cells 30 (2012), pp. 2843–2847.

[13] Pascale Bouchard-Cannon et al. “The circadian molecular clock regulates adult hippocampal neurogenesis by controlling the timing of cell-cycle entry and exit”. In: Cell reports 5.4 (2013), pp. 961–973.

[14] Maria P Alcolea et al. “Differentiation imbalance in single oesophageal progenitor cells causes clonal immortalization and field change”. In: Nature cell biology 16.6 (2014), p. 615.

[15] A. M. Baker et al. “Quantification of crypt and stem cell evolution in the normal and neoplastic human colon”. In: Cell Rep 8.4 (Aug. 2014), pp. 940–947.

[16] Samira Chabab et al. “Uncovering the Number and Clonal Dynamics of Mesp1 Progenitors during Heart Morphogenesis”. In: Cell reports 14.1 (2016), pp. 1–10.

[17] Michael Flossdorf et al. “CD8+ T cell diversification by asymmetric cell division”. In: Nature immunology 16.9 (2015), pp. 891–893.

[18] Julie K Watson et al. “Clonal dynamics reveal two distinct populations of basal cells in slow-turnover airway epithelium”. In: Cell reports 12.1 (2015), pp. 90–101.

[19] A. Marciniak-Czochra et al. “Modeling of asymmetric cell division in hematopoietic stem cells-regulation of self-renewal is essential for efficient repopulation”. In: Stem Cells Dev. 18 (2009), pp. 377–385.

[20] Daniel T Gillespie. “Exact stochastic simulation of coupled chemical reactions”. In: The journal of physical chemistry 81.25 (1977), pp. 2340–2361.

[21] Frederik Ziebell, Ana Martin-Villalba, and Anna Marciniak-Czochra. “Mathematical modelling of adult hippocampal neurogenesis: effects of altered stem cell dynamics on cell counts and bromodeoxyuridine-labelled cells”. In: Journal of The Royal Society Interface 11.94 (2014).

[22] Joseph Magill and Jean Galy. Radioactivity, radionuclides, radiation. Springer, 2005.

[23] Kenneth P Burnham and David R Anderson. Model selection and multimodel inference: a practical information-theoretic approach. Springer, 2002.

[24] Qiuhao Qu et al. “Orphan nuclear receptor TLX activates Wnt/*β*-catenin signalling to stimulate neural stem cell proliferation and self-renewal”. In: Nature cell biology 12.1 (2010), pp. 31–40.

[25] Roeben N Munji et al. “Wnt signaling regulates neuronal differentiation of cortical intermediate progenitors”. In: The Journal of Neuroscience 31.5 (2011), pp. 16761687.

[26] B Li et al. “Multitype Bellman-Harris branching model provides biological predictors of early stages of adult hippocampal neurogenesis”. In: BMC Systems Biology to appear (2017).

[27] Karl Popper. The logic of scientific discovery. Hutchinson, 1959.

[28] Xiang Gao et al. “Conditional knock-out of *β*-catenm in postnatal-born dentate gyrus granule neurons results in dendritic malformation”. In: Journal of Neuroscience 27.52 (2007), pp. 14317–14325.

[29] D. C. Lie et al. “Wnt signalling regulates adult hippocampal neurogenesis”. In: Nature 437.7063 (Oct. 2005), pp. 1370–1375.

[30] Theologos M Michaelidis and D Chichung Lie. “Wnt signaling and neural stem cells: caught in the Wnt web”. In: Cell and tissue research 331.1 (2008), pp. 193210.

[31] Heiko Enderling et al. “A mathematical model of breast cancer development, local treatment and recurrence”. In: Journal of theoretical biology 246.2 (2007), pp. 245–259.

[32] T. Stiehl and A. Marciniak-Czochra. “Characterization of stem cells using mathematical models of multistage cell lineages”. In: Mathematical and Computer Modelling 53 (2011), pp. 1505–1517. ISSN: 0895-7177.

[33] Suzanne L Weekes et al. “A multicompartment mathematical model of cancer stem cell-driven tumor growth dynamics”. In: Bulletin of mathematical biology 76.7 (2014), pp. 1762–1782.

[34] Allon M Klein et al. “Mouse germ line stem cells undergo rapid and stochastic turnover”. In: Cell Stem Cell 7.2 (2010), pp. 214–224.

[35] David P Doupé et al. “A single progenitor population switches behavior to maintain and repair esophageal epithelium”. In: Science 337.6098 (2012), pp. 1091–1093.

[36] Katrin Busch et al. “Fundamental properties of unperturbed haematopoiesis from stem cells in vivo”. In: Nature 518.7540 (2015), pp. 542–546.

[37] Franziska Michor et al. “Dynamics of chronic myeloid leukaemia”. In: Nature 435.7046 (2005), pp. 1267–1270.

[38] N. Hayes and RS Nowakowski. “Dynamics of cell proliferation in the adult dentate gyrus of two inbred strains of mice”. In: Developmental brain research 134.1 (2002), pp. 77–85.

[39] Stefano Farioli-Vecchioli et al. “Running rescues defective adult neurogenesis by shortening the length of the cell cycle of neural stem and progenitor cells”. In: Stem Cells 32.7 (2014), pp. 1968–1982.

